# Automated Machine Learning Profiling with MAP-HR for Quantifying Homologous Recombination Foci in Patient Samples

**DOI:** 10.1101/2025.02.20.639329

**Authors:** Tugba Y. Ozmen, Matthew J. Rames, Gabriel M. Zangirolani, Furkan Ozmen, Gordon B. Mills

## Abstract

Accurate visualization and quantification of homologous recombination (HR)-associated foci in readily available patient samples are critical for identifying patients with HR deficiency (HRD) when they present for care to guide polyADP ribose polymerase (PARP) inhibitors (PARPi) or platinum-based therapies. Immunofluorescence (IF) assays have the potential to accurately visualize DNA repair processes as punctate foci within the nucleus. To ensure precise HRD assessment, we developed MAP-HR, (**M**achine-learning **A**ssisted **P**rofiling of **H**omologous **R**ecombination), a scalable machine-learning (ML) analysis platform to enable effective patient triage and therapeutic decision-making. This workflow integrates high-resolution four-channel IF imaging and automated analysis of Geminin (cell cycle states), RAD51 (HR repair), *γ*H2AX foci (double strand breaks) and DAPI (nuclear localization) in both cultured cell lines and in a single formalin-fixed, paraffin-embedded (FFPE) patient sample. Using a spinning disk confocal microscope, we optimized imaging parameters to improve resolution and signal-to-noise ratio. Our MAP-HR pipeline uses nested nuclei and segmentation of foci to analyze the HR status of each cell, unlike competing bulk or single-foci marker assays, allowing evaluation of HR functional heterogeneity across and within patient biopsies. This approach facilitates robust comparisons of HR and foci-based processes across diverse cell populations and patient tissues, enabling scalable, translational research.

**GRAPHICAL ABSTRACT:** 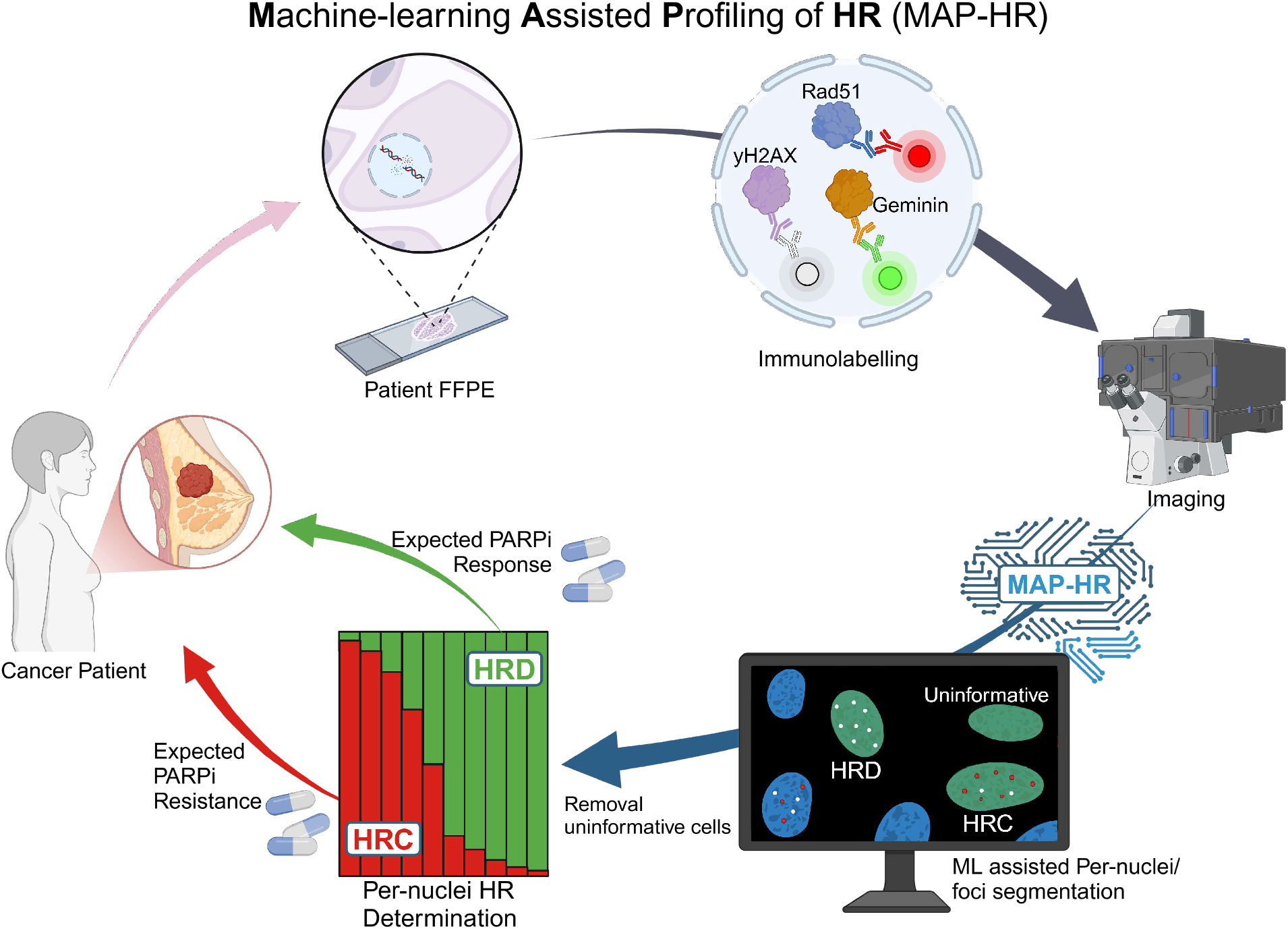

## INTRODUCTION

Accurate DNA replication and repair are essential for maintaining genomic integrity and preventing deleterious consequences such as mutations and chromosomal instability. In cancer, defects in DNA repair pathways are common and contribute significantly to tumor initiation, progression, and resistance to therapy^1^. Among DNA damage types, double-strand breaks (DSBs) are particularly toxic and require precise repair mechanisms to maintain genomic stability^2^. Homologous recombination (HR) is the preferred high-fidelity repair pathway that utilizes a sister chromatid as a template for error-free repair of DSBs during the G2 phase of the cell cycle.^3^ During this process, H2AX is rapidly phosphorylated, forming *γ*H2AX^4^. This phosphorylation marks damage sites and helps recruit repair proteins including RAD51, a key protein in HR repair, that are loaded onto the break site^5^. HR deficiency (HRD) has been linked primarily to mutations in BRCA1/2, PALB2 and RAD51 allowing genetic screening to be used to help triage patients with these biomarkers for treatment with poly ADP ribose polymerase (PARP) inhibitors (PARPi) that are synthetically lethal with HRD^6,7^. However, preclinical and clinical data in multiple cancer types indicate that BRCA1/2 wild type (*wt*) tumors can also exhibit HRD phenotypes^8–10^, which has been demonstrated to render them susceptible to treatment with PARPi or platinum-based treatments due to their inability to efficiently repair DSBs induced by therapy. Immunofluorescence-based assays have emerged as valuable tools for visualizing active repair processes, which appear as punctate foci within the nucleus^11,12^. To ensure accurate HRD detection, a robust and scalable analysis platform is essential for effective patient triage and informed therapeutic decision-making. Despite the successful integration of several software solutions for cell and nuclei segmentation^13,14^, most do not provide reliable segmentation of intranuclear foci. This gap highlights the urgent need for specialized, custom segmentation pipelines capable of addressing these challenges to select patients who are most likely to benefit from HR-targeted therapies.

Foci analysis of HR markers has emerged as a powerful method for evaluating functional HR processes offering a distinct advantage over genomic scar assays that reflect a history of HRD^15^, by providing real-time insights into the current HR status in a cell or tumor. Many foundational assays in this field have relied upon manual foci annotation for the basis of functional HR calls^16,17^, spurring the need for automated analysis which can mitigate biases and improve throughput. Although upstream nuclear segmentation pipelines are fairly robust and commonplace in IF-based applications^14,18,19^ integrated foci-specific segmentation and analysis requires further development. Common problems arise due to variations in: nuclear background, small foci size, focal deviations, poor signal-to-noise, batch variation, and intra- and inter-tumoral variation^20^. Traditional foci-based analyses typically leverage local noise reduction filters which attempt to enhance local signal from putative foci^21–23^. Secondarily thresholding methods are often used to segment foci via their peak intensities^24^. While these methods have dramatically improved abilities to extract relevant foci from input datasets, they still encounter difficulties in: large batch variations, normalizations, and accurate foci size detection. Recent commercial software packages like Zeiss Arivis have begun integrating foci-based feature extractions, yet generally operate on the same fundamental principles as previous platforms.

Motivated by growing unmet clinical needs, here we present MAP-HR (**M**achine-learning **A**ssisted **P**rofiling of **H**omologous **R**ecombination), a semi-automated multi-level ML pipeline to extract and interpret functional foci across clinical cohorts. To better account for differences in foci presentation due to batch effects, labeling differences, background expression, our MAP-HR model training focuses on foci shape rather than solely intensity. By not relying on thresholding for final foci quantifications, MAP-HR lends itself to large-scale downstream analysis^8,25^ and thresholding via R. MAP-HR offers enhanced flexibility for downstream data curation and normalization, critical foundational steps behind any clinical-facing assay.

## MATERIALS AND METHODS

### Sample Collection

This study was approved by the Oregon Health & Science University (OHSU) Institutional Review Board (IRB). All biospecimens were collected under the MMTERT observational study (Mitri 2018) (IRB #16113).

### Cell culture

UWB1.289 and UWB1.289+BRCA1 cell lines were obtained from ATCC (Manassas, VA, catalog #CRL-2945, #CRL-2946). The UWB1.289 cells were cultured in 50% RPMI Medium (Gibco, catalog #1640) and 50% MEGM Bullet Kit (Lonza, catalog #CC-3150) with 3% (v/v) FBS (Gibco, catalog #16140-089). UWB1.289 cells were cultured in the same base media mixture with the inclusion of Geneticin (Gibco, catalog #10131035). Cells were seeded at 4×10^4^ cells per well in 8-well coverslide (Cellvis, catalog #C8-1.5H-N). After 24 hrs, cells were exposed to varying Gemcitabine (Selleck chemicals, catalog #s1714) concentrations for an additional 24 hrs.

Resulting treated cells were fixed with 4% paraformaldehyde (Thermo Fisher, catalog #28908), and subjected to IF.

### Immunofluorescence Staining and Imaging

For IF staining of formalin-fixed, paraffin-embedded (FFPE) tissue samples, 3 µm sections were deparaffinized using xylene twice for 10 minutes, 100% ethanol twice for 10 minutes, 95% ethanol for 5 minutes, 70% ethanol for 5 minutes, and 50% ethanol for 5 minutes, and then left in PBS. Tissues were antigen retrieved in a pressure cooker using Citrate (pH 6) and DAKO antigen retrieval (pH 9) buffers (Sigma, catalog #c9999). and Agilent, catalog #s236784-2, respectively). Tissues were washed twice in PBS and then were further permeabilized with 0.4% Triton X-100 in PBS for 1 hour. The excess permeabilization solution was removed by washing three times in PBS, and then a hydrophobic barrier was placed around the mounted tissue.

Tissues were blocked with 3% BSA (Fisher BP1600-100 in PBS prior to antibody labeling. The tissues were then incubated with primary antibodies RAD51 (Abcam, catalog #ab133534), Geminin (Leica, catalog #50-255-2243), *γ*H2AX (Sigma, catalog #05-636) on a rocker for 2 hours at room temperature in a humidity chamber. After three PBS washes, secondary antibodies were applied: CF-680-conjugated goat anti-mouse(IgG1) (Biotium, catalog #20253) and Alexa Fluor 555-conjugated goat anti-rabbit (Thermo Fisher, catalog #a21428). Alexa Fluor 488-conjugated goat anti-mouse (IgG2a) (Thermo Fisher, catalog #A-21131). After three final PBS washes, all tissue samples were postfixed with 4% PFA for 10 minutes at room temperature. Hoechst staining solution (1:5000) was used and mounted in Prolong gold antifade (Invitrogen, catalog #P36930. Sample images were acquired using a spinning disk confocal microscope with a 40x objective. A minimum of 100 tumor cells per sample were used for downstream analysis.

### Imaging optimization and orthogonal projections

In order to assess imaging-specific and magnification optimizations, a representative TNBC patient was stained as outlined above and imaged with both 20x Zeiss Axioscan7 and 40x Zeiss-Yokogawa CSU-X1 Spinning disc confocal microscopes. Line profiles from 100 foci were manually drawn to avoid multiple foci across 3 µm lateral segments for each foci marker. These lines were drawn at the 5^th^ Z-slice from the 3 µm vertical stacks (300 nm increments) as acquired from the spinning disc confocal microscope. Downstream organization of extracted line profiles from *Fiji* (saved as .*csv* files) were organized in *R Studio* to plot and compare FWHM from observed foci intensity profiles. 2D Averaging was performed by taking the average intensity across each lateral view, while the Z-profile was taken from the vertical intensity profiles at the peak foci position as found from 2D line profiles. Subsequent plots compare the performance of the same 100 manually annotated foci, when compared by lateral, projection, or axial views from the raw data.

### Model training and segmentation performance

Nuclear and foci segmentation models were trained from manual annotations using orthogonal cell/FFPE patient datasets. Input annotations from a collection of foci positive and foci negative samples were used for model training. In particular, a focus on providing a variety of nuclear morphologies and intensities during model training was used to improve segmentation model flexibility. Despite this, batch-specific scaling factors can be used to boost or dimmish DAPI intensities to aid in nuclear segmentations. Foci segmentation was trained on a variety of foci intensities, within both foci positive and known foci negative samples, by focusing on punctate features across a wide range of intensities, downstream thresholding and batch corrections should be more readily applicable. Foci segmentation performance in particular was performed by comparing manual annotation of *γ*H2AX and RAD51 foci within 100 representative nuclei from a TNBC with known HR-competency.

### Image segmentation and feature extraction

Raw image files from *Zen* (.*czi*) were imported into custom *FIJI*^26^ macro software using bio-formats^27^ for nuclear and foci segmentation. A schematic overview of MAP-HR can be seen in (**Supplementary Figure 3**). Input images were first split into their respective channels for: 1) DAPI, 2) *γ*H2AX foci 3) RAD51 foci, and 4) Geminin cell phase nuclear staining. Using split nuclei channels, each sample utilizes pre-drawn tumor .*roi* files to run nuclear segmentation models through *WEKA*. Binarized nuclear output images are then subjected to post-segmentation nuclei cleanup through *dilate, smoothing, fill gaps*, and *erode* functions within *Fiji*. Adjacent nuclei are subsequently subdivided using watershedding, wherein nuclei > 3 µm^2^ were saved as .*roi* files for downstream foci segmentation. These same nuclear boundary .*roi* files were subsequently applied to the Geminin channel to extract total geminin nuclear staining per cell for use in later cell-phase identifications. Extracted total nuclear positions and marker expressions were saved in Nuclear Feature extraction tables (.*csv*).

After nuclear and geminin feature extraction, each foci channel (*γ*H2AX and RAD51) was separately loaded with corresponding pre-trained foci segmentation models in *WEKA*. Within each nuclear. roi from upstream nuclear segmentation, outside signal was removed, and foci segmentation was performed on nuclei with average foci marker expression above the global background. This additional step improves total analysis run-times by skipping unnecessary segmentation on nuclei without observable foci. After *WEKA* segmentation for marker-specific foci, output foci feature tables (.*csv*) are extracted per nucleus and run analogously on all nuclei from upstream nuclear segmentation. Aggregate feature tables from both nuclei and matching foci where available (.*csv*) ready for downstream analysis.

### Data organization, batch corrections, thresholding, and data visualization

Full output feature tables saved from all samples were loaded into custom post-processing script written in *R*. Utilizing unique nuclear identifiers from both nuclear and foci level segmentation results, foci segmentation output data frames were paired with matching parental nuclei. Additionally, using a global .*roi* reference .*csv* per patient, each nuclei position and foci was corrected back to its corresponding global position. Since *Fiji* processing considers each smaller image using a new coordinate system, global positions must be offset after the nested segmentation and feature extraction on smaller nuclear features is performed.

If utilizing shared reference samples within multiple cohorts, the next layer allows assigning references for *log(mean)* normalization scaling factors based on shared reference samples. For optimal performance, we run both HRC and HRD positive and negative control biopsies as references across cohorts. After applying batch corrections, nuclear and foci-specific filters are applied. Critically, utilizing HRC and HRD references ensures both positive and negative foci intensity thresholds for downstream nuclear calls.

Nuclear HRC and HRD calls were performed utilizing decision tree outlined in Figure 1b. For finding geminin positivity, G2 thresholds were calculated from bimodal intercept of Gaussian distributions (See also Supplemental Figure 5). This type of gaussian mixture model run per sample helps to identify levels of Geminin positivity within sample and potentially lineage-specific contexts. This threshold ensured selection of all late S/G2 cells per sample. Within G2 nuclei, foci minimum intensity thresholds were set using HRD/untreated controls which lacked sufficiently bright foci. Foci nuclear positivities were then called as follows: RAD51 positivity was defined as a minimum of 5 foci greater than 500 nm in diameter, and *γ*H2AX positivity was defined as a minimum of 2 foci greater than 300 nm in diameter. After nuclei calls were found per sample, sample sets/cohorts are plotted in tandem using *ggplot2* package in *R*, wherein *“fill”* scaling is used to see the relative proportion of each foci positivity call. Samples with less than 100 Geminin+ nuclei were called unquantifiable, as visualization of limited G2 populations hinders reliable detection of cells actively performing HR/encountering HRD.

**Figure 1:**
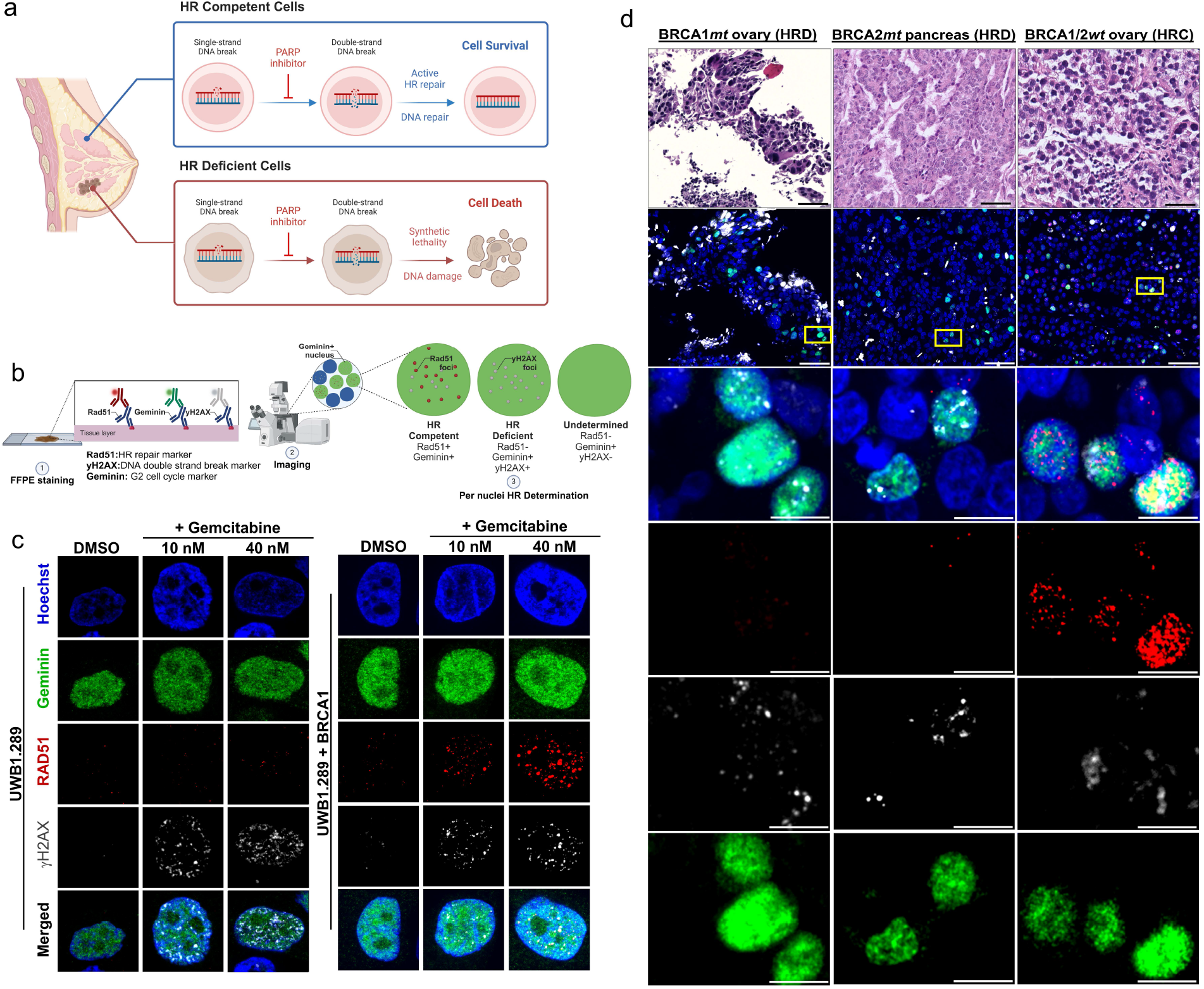
Characterizing Homologous Recombination Deficiency in Cancer Cell Line Models and Patient Tissues. a) Representative illustration of synthetic lethality linked to Homologous Recombination (HR) Competency vs Deficiency. Created in BioRender. Yildiran ozmen, T. (2025) https://BioRender.com/z60f958b) HR competency and deficiency decision tree via immunofluorescence assay and foci-based analysis. c) Representative UWB and UWB+BRCA1 restored cells treated with Gemcitabine. Cell nuclear insets are 5 µm wide. d) Representative H&E, and functional immunofluorescence imaging of RAD51/yH2AX from patient FFPE tumors with known BRCA status classified as HR deficient (HRD) and HR competent (HRC). Scale bars are 50 µm and 10 µm for full and magnified regions respectively.

## RESULTS

### Visualizing Homologous Recombination in Cells and Patient Samples

HR is a high-fidelity DNA repair mechanism and the preferred pathway for repairing DSBs without the introduction of genomic errors. PARPi induce synthetic lethality in HRD cells by inhibiting single-strand break (SSB) repair, thereby generating DSBs through subsequent cell cycles and increasing the cell’s reliance on HR repair^28^ **(Figure 1a)**. To determine which cells are actively performing HR, we implemented a decision tree that stratifies cells based on HR competency versus deficiency, assessed solely in the G2 phase (where functional HR occurs) as identified by Geminin-positive staining **(Figure 1b)**. Cells actively performing HR are identified by the presence of >5 RAD51 foci in G2 cells and are termed HR-competent (HRC). In contrast, cells with unresolved DSBs in G2, marked by >2 *γ*H2AX foci, and lacking active RAD51 foci are classified as HR-deficient (HRD). Notably, G2 phase cells without RAD51 or *γ*H2AX foci are considered “Undetermined” because HR only occurs when DSBs are present, and the absence of repair (RAD51) or marked damage (*γ*H2AX) makes it unclear what the potential HR capacity is for that cell.

To illustrate the HR process in cells, we conducted IF staining for the three markers (RAD51, *γ*H2AX, Geminin) and DAPI for nuclear localization in both BRCA1-deficient and in BRCA1-restored UWB cells **(Figure 1c)**. Upon treatment with the DNA-damaging agent Gemcitabine, both cell lines exhibited clear *γ*H2AX (DSB) foci formation; however, only the BRCA1-restored UWB cells demonstrated functional RAD51 (HR repair) foci. Importantly, these same markers also reflect similar processes in patient formalin-fixed, paraffin-embedded (FFPE) samples **(Figure 1d)**. Representative H&E staining paired IF imaging, and magnified fields of view (FOV) reveal that BRCA1 and BRCA2 mutant (*mt*) tumors lack sufficient RAD51 foci, while unresolved *γ*H2AX DSBs are visible in G2-phase tumor cells. In contrast, BRCA1/2 *wt* tumors display HR competency in G2 phase cells through RAD51 foci formation. While this process can be visually observed in patient biopsies, we next developed an analytical method to quantitatively extract these features to allow determination of status of the patient tumor for research and eventual clinical application.

### MAP-HR Segmentation and Analysis Workflow

The integration of foci-level details across diverse aspects such as tumor-rich regions, nuclear morphology and positioning, bulk nuclear expression, and the specific positioning and size of each foci is essential to quantify DNA repair capacity. Most competing methods, however, tend to focus on only subsets of these factors, often isolating bulk or individual marker level details without accounting for their interplay. To enhance foci quantification for patient biopsies, we developed a scalable framework that integrates spatial, morphological, and expression data across tumor-rich regions of interest (ROIs) that can organize these details (**Supplemental Figure 1**). Tumor ROIs are selected, and geminin expression is analyzed to determine the cell cycle phase. Using image-offset spatial coordinates, we map back each corrected ROI and extract bulk nuclear intensities including Geminin intensities for G2 stratification **(Supplemental Figure 2&3)**. Nuclei segmentation within ROIs provides nuclear-level data, including position, diameter, and morphology, while foci-level parameters such as position, intensity, diameter, and total foci count are quantified for each nucleus. This multiscale approach feeds into a decision tree model to stratify co-occurring HR-related processes. By aggregating foci attributes across patient cohorts and reference samples, our method identifies patterns in HR repair activity at tumor rich areas, nuclear, and foci scales, enabling robust biomarker discovery and improved patient stratification.

**Figure 2:**
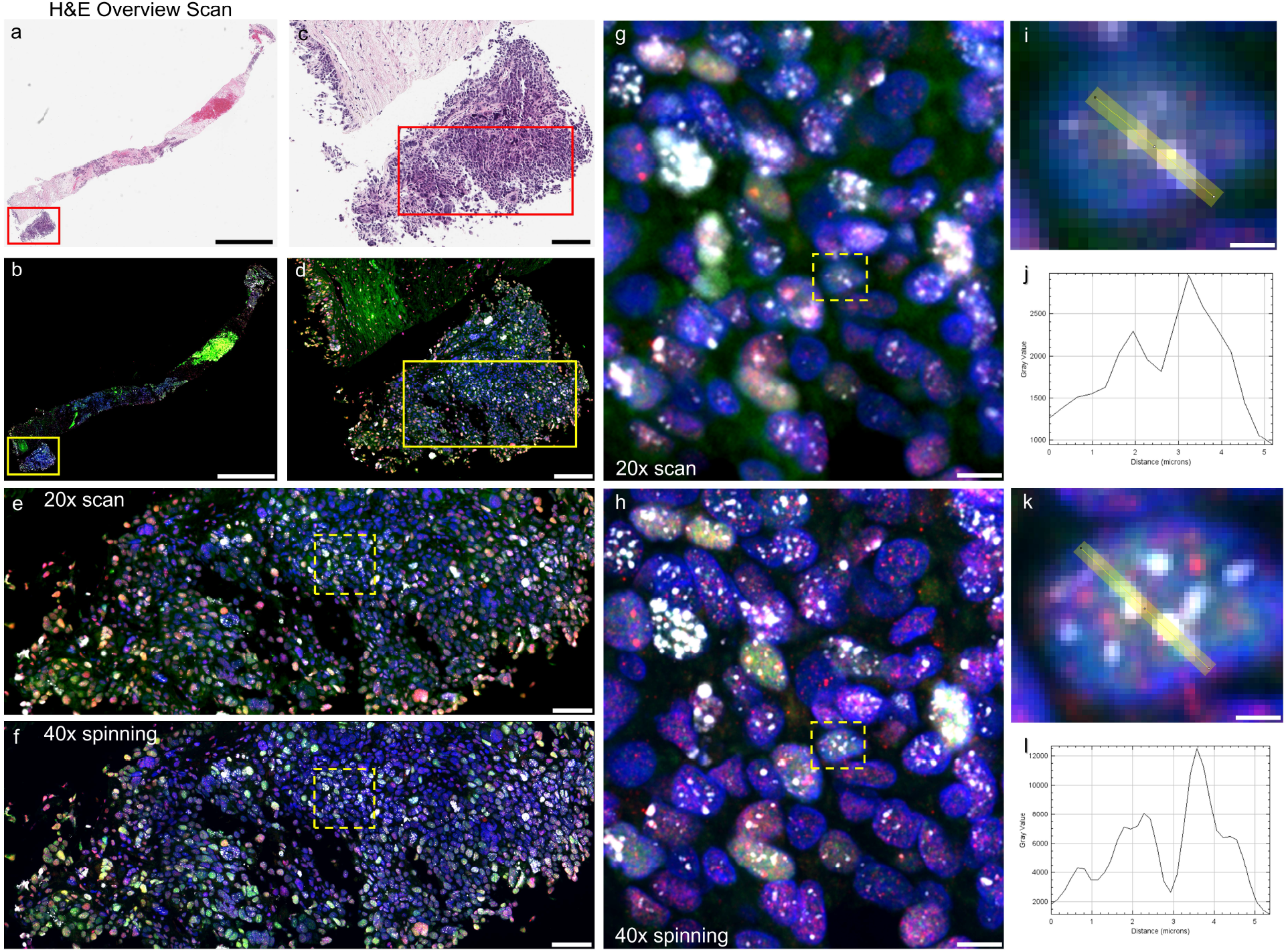
Improving Imaging Quality through Microscope Optimization. a) H&E overview scan from breast cancer core biopsy. Pathologist annotated tumor rich areas are shown in red box. Scale bar 500 µm. b) Identical FOV from adjacent section with nuclear staining via hoechst immunolabeled for yH2AX (white), RAD51 (red), and Geminin (green) scanned with 20x objective. Scale bar 500 µm. c) Pathologist annotated tumor-rich area selection from breast cancer core biopsy. Scale bar 100 µm. d) Corresponding area from c as scanned from b. Scale bar 100 µm. e) Magnified tumor-rich region from d as indicated in yellow box using 20x scanner. Scale bar 30 µm. f) Corresponding area from e as imaged with a 40x spinning disk confocal microscope. Scale bar 30 µm. g) Magnified ROI from e as indicated by dashed yellow box. Scale bar 5 µm. h) Magnified ROI from f as indicated by dashed yellow box. Scale bar 5 µm. i) Representative nuclei with marked line profile across multiple yH2AX foci as seen from g. Scale bar 1 µm. j) Plotted line profile of yH2AX from i. (average line width=2 pixels) k) Identical representative nuclei with marked line profile across multiple yH2AX foci as seen from h. Scale bar 1 µm. l) Plotted line profile of yH2AX from k. (average line width=3 pixels)

### Foci imaging Optimization

As the backbone behind any functional punctate analysis, imaging optimizations which retain optimal resolution to accurately discern foci size, shape and intensity is critical. To exemplify these optimizations in raw imaging quality, we immunolabeled a triple negative breast cancer (TNBC) tumor for: 1) Cell phase via Geminin, 2) HR repair via RAD51, and 3) DSBs via *γ*H2AX, using H&E to guide imaging for tumor-rich regions (**Figure 2a-d**). Next, we compared imaging using 20x scanning or 40x spinning disk confocal (See methods) (**Figure 2e & f**), wherein despite the ability to visualize foci through both microscopes, the 40x spinning disk confocal revealed substantially crisper, better resolved and more numerous foci within tumor cells (**Figure 2g & h**). To better visualize differences in raw foci resolution and signal-to-noise (SNR) contrast, we compared line-profiles within the same nucleus for *γ*H2AX foci as observed with both microscopes (**Figure 2i-l**). 20x scanning appeared to reveal 2 foci with SNR ~1.5-2 and diameters of nearly a micron, whereas 40x spinning disc, due in part to limited out of focus illumination and higher lateral resolution, yielded more foci with SNR ~2-6 and diameters closer to 600 nm by Full-Width-Half-Maximum (FWHM) fitting.

Next, to account for the 3D nature of these HR-related protein complexes, we quantified the relative 2D and 3D resolutions seen via 2D or 3D Z-stack image acquisitions. Considering patient FFPE sections are commonly sectioned at 4-5 µm thickness, we collected 3 µm thick Z-stacks at 300 nm intervals to characterize foci seen via 3D within the biopsy (**Figure 3a**).

**Figure 3:**
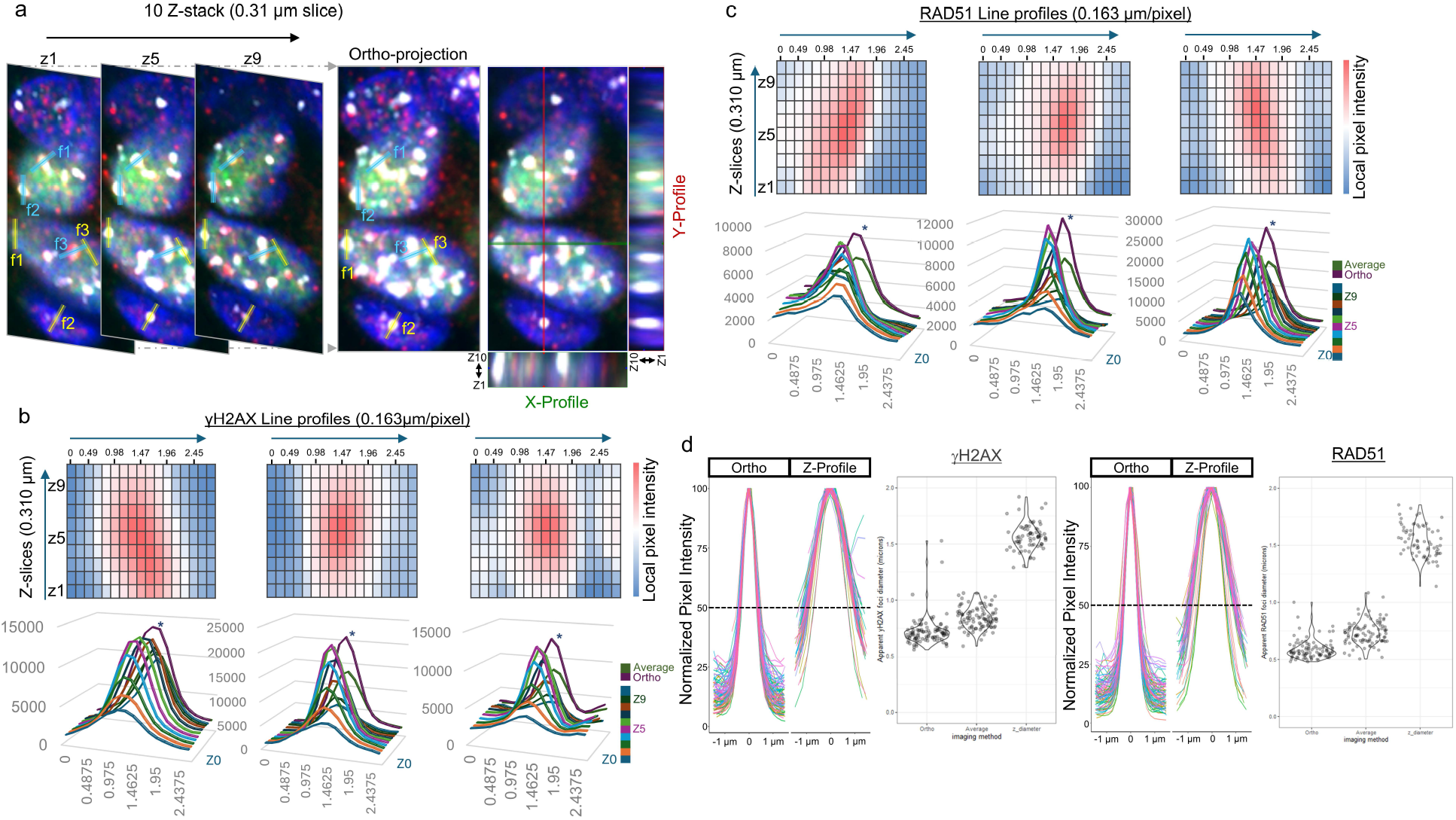
Refining Image Capture to Enhance Analytical Precision. a. Representative Z slices from FFPE with nuclear staining via hoechst immunolabeled for yH2AX (white), RAD51 (red), and Geminin (green) taken with 40x confocal at 0.31 um per Z and matching orthogonal projection b. Selected yH2AX line profiles and corresponding intensities along each Z slice. 3D visualization of intensity profiles along each Z-slice paired to orthogonal projection intensity profile indicated via *. c. Selected RAD51 line profiles and corresponding intensities along each Z slice. 3D visualization of intensity profiles along each Z-slice paired to orthogonal projection intensity profile indicated via *. d. Relative distribution of apparent XY diameters vs Z depth for 100 yH2AX & RAD51 foci calculated using full-width half maximum along either orthogonal or Z-profiles per matching foci.

Numerous RAD51 and *γ*H2AX foci can be seen across each section, which can be vertically compressed into a maximum intensity orthogonal projection retaining each foci seen in 3D as a 2D representation. Importantly, as seen via X/Y axis profiles with Z, the apparent lateral resolution is substantially better than the axial resolution, a well-known aspect in fluorescent imaging applications^29^. To better visualize the relative differences in 2D, 3D, and orthogonal projections, we plotted the lateral and axial line profiles of 3 representative *γ*H2AX and RAD51 foci (**Figure 3b & c**). While the apparent foci intensity profiles vary per Z-slice, the orthogonal projection profile retains the properties of the peak Z-slice, visually outperforming any average view seen through the 3 µm Z height. Extending this style of line-profiles to 100 *γ*H2AX and 100 RAD51 foci, normalized intensities further demonstrate the deviations in FWHM resolution by average lateral resolution via a maximum intensity orthogonal projection (Ortho) compared to the average axial foci resolution (Z-profile) (**Figure 3d**). Orthogonal projections of *γ*H2AX and RAD51 foci were typically ~600 nm, outperforming both Averaged 2D views and Axial FWHMs at ~ 700-800 nm and ~1.6 microns respectively. Collectively, this analysis demonstrates that collecting 3D image stacks, and then vertically compiling them through an orthogonal projection, retains the most precise resolution for accurate foci quantification. This improvement in imaging quality requires slightly increased image acquisition times for 3D Z-stacks, yet downstream processing utilizing the 2D maximum intensity orthogonal projection should greatly improve computing time over 3D methods.

### Multi-level Semi-automated Machine Learning Foci Extraction Pipeline

To develop a more scalable foci quantification tool for use in analyzing patient biopsies, we first outlined how data should be organized. Aggregate foci attributes across patient cohorts and known reference samples could be utilized to stratify HR competency. To encapsulate these details, we implemented a nested approach which sought to pair higher level patient and tumor positional data alongside per-nuclear protein expression and sub-nuclear foci specific properties (**Figure 4a**). To increase accessibility, we developed MAP-HR pipeline using the FIJI(*ImageJ)*^30^ macro-language, that can automate the processes from image segmentation to feature extraction and provide organized data (**Supplemental Figure 3a**). While each function in *FIJI* is typically performed within smaller ROI subsets (i.e tissue ROIs down to smaller foci ROIs), positional offsets which can realign multiple regions back onto their global positions(**Supplemental Figure 2**) from higher level nested ROIs can ensure accurate positioning for each level of detail (smaller tumor ROI regions) within downstream analyses, such as through *R* (**Supplemental Figure 3b**).

**Figure 4:**
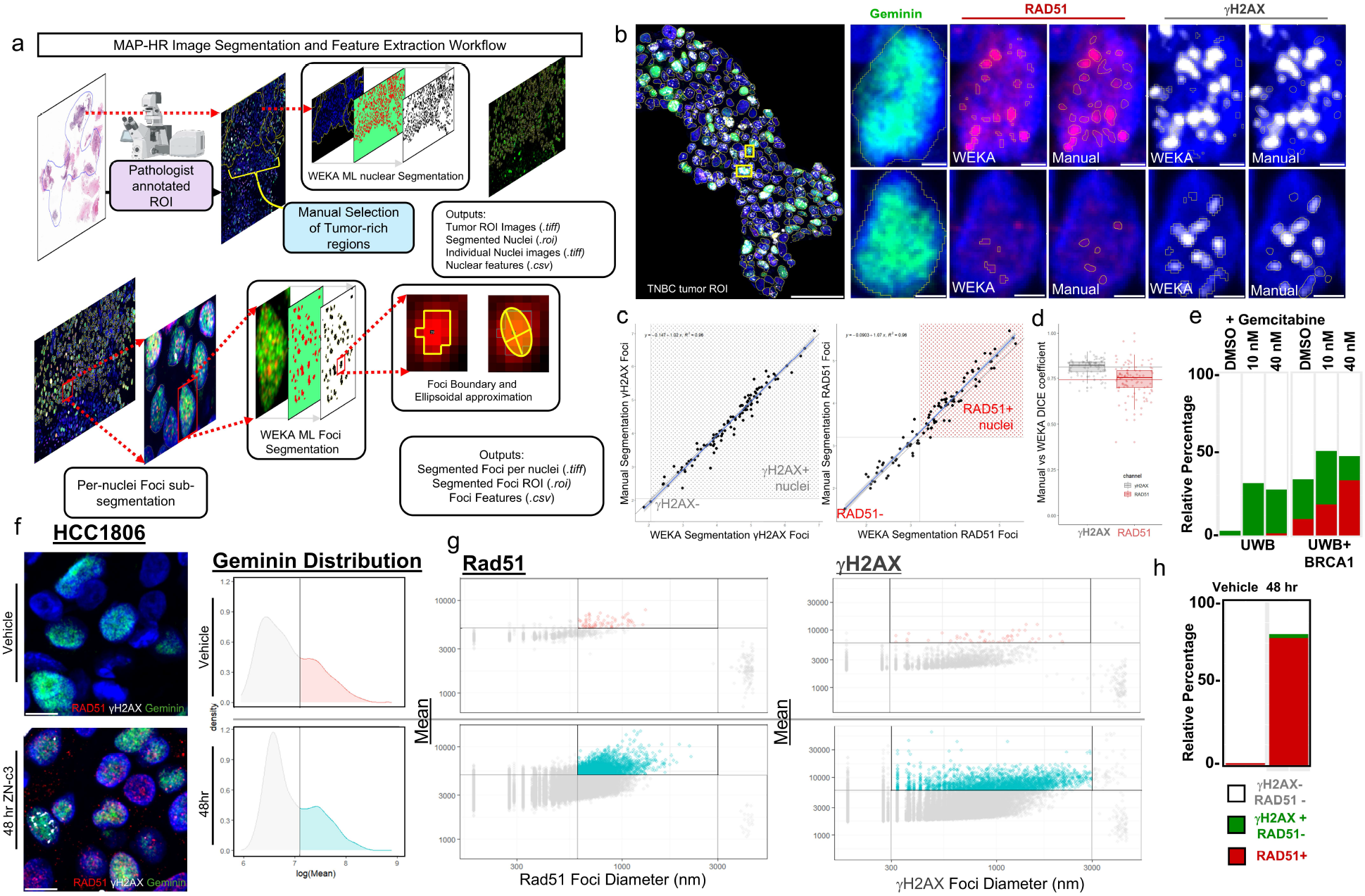
MAP-HR Image Segmentation and Feature Extraction Workflow and Performance. a) Image segmentation and feature extraction using MAP-HR pipeline utilizing nuclei and sub-nuclear foci segmentation via nested machine learning analysis. b) Representative FOV from TNBC patient IF imaging of RAD51/yH2AX/Geminin staining and example foci segmentation via manual or WEKA ML outputs. Scale bars 50 µm and 2 µm in each inset c) Correlation plots for both manual and WEKA ML foci segmentation when convolving foci diameter, intensity, and number detected from 100 representative nuclei. Convolved thresholds reveal analysis outputs for each respective foci positivity by either manual or WEKA segmentation outputs.d) DICE coefficients for both yH2AX and RAD51 were calculated to be 0.8 and 0.74 respectively. DICE is calculated as 2 times overlapping area over the cumulative total area from both manual and WEKA segmentation outputs. e) Relative percentage of HRC/HRD/Uninformative nuclei populations for both UWB and UWB+BRCA1 cell populations when treated with increasing dosages of Gemcitabine. f) HCC1806 xenograph images with/without ZN-c3 treatments. Scale bars 10 µm. Additional Geminin intensity profile distributions when plotted on log-scale, with threshold for Geminin positivities filled as indicated on the right-side of each distribution for both vehicle and ZN-c3 treated samples colored pink and blue respectively. g) Dot-plot of RAD51 and yH2AX foci when plotted by foci diameter and foci mean intensity both on log-scales. Regions thresholded and colored pink and blue respectively indicate positive foci per marker. h) Relative percentage of HRC (RAD51+, Geminin+) / HRD (RAD51-, yH2AX+, Geminin+) / Uninformative (RAD51-, yH2AX-, Geminin+) nuclei populations from f, after applying displayed foci thresholds shown in g.

### Machine Learning Foci Segmentation Performance

To validate model performance, we imaged a representative FOV from a TNBC patient for Geminin, RAD51, and *γ*H2AX (**Figure 4b**). Through manual annotation compared to WEKA ML outputs, we can see good concordance between selected foci and sizes (**Figure 4b**). By convolving the relative signal between foci size, intensity, and number, we plotted the correlation between manual vs WEKA performance (**Figure 4c**), with R^2^ value > 0.96 for both models. Each nucleus foci convolved score (for purposes of checking segmentation performance) was calculated using: 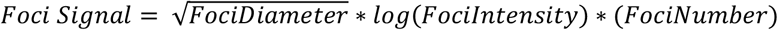. Individual attribute correlations were also found for *FociDiameter, FociIntensity*, and *FociNumber* for each marker respectively (**Supplemental Figure 4**). Additionally, to show the degree of concordance between manual and machine learning segmentation outputs, we calculated Dice coefficients for both foci and found performance per foci were 80% for *γ*H2AX and 74% for RAD51 (**Figure 4d**). In order to demonstrate the ability to quantify increases in RAD51 and *γ*H2AX foci formation through MAP-HR, we extracted the formation of HRC and HRD nuclear populations from UWB and UWB+BRCA1 restored cell lines as shown from Figure 1c (**Figure 4e**). Through normalizing the number of G2 phase cells for each treatment condition, we observed an increase of *γ*H2AX foci + nuclei in HRD UWB cells induced by Gemcitabine treatment. Additionally, BRCA1 restored UWB cells demonstrated a clear increase in HRC cell populations which were actively performing HR in response to DSBs induced by Gemcitabine treatment. In the absence of Gemcitabine treatment in UWB+BRCA1 G2+ cells, methods that define HRD based solely on RAD51-negative and Geminin-positive status, without simultaneously assessing *γ*H2AX, may incorrectly classify UWB+BRCA1 cells as HRD leading to the misinterpretation of HRC cell line as HRD.

### Representative Foci Thresholding and Downstream Analytics in R

After validating model performance, we tested MAP-HR on a representative set of cell line xenographs (HCC1806) comparing both vehicle and 48 hr treatment with the WEE1 inhibitor azenosertib (ZN-c3) that inhibits the S and G2 phase checkpoints driven by the ATR/CHK1/WEE1 cascade ^25^ (**Figure 4f)**. As seen in raw images, azenosertib treatment increased DSBs indicated by the presence of *γ*H2AX foci in cells with RAD51 foci representing ongoing HR repair. Once again in the absence of azenosertib, there were insufficient *γ*H2AX foci + nuclei to accurately assess HR competence. To ensure correct assignment of HR, we additionally filtered by cell phase using geminin intensity positivity through gaussian distribution profiles (**Supplemental Figure 5**). Through modeling the 2 most prominent gaussian distributions, we calculated the intercept and utilized it as the putative G2 phase cutoff per sample. Using vehicle controls as references, dot-plots of RAD51 and *γ*H2AX foci diameter and intensities were thresholded to retain substantially large and bright foci as functional foci (**Figure 4g**). Foci diameters for *γ*H2AX and RAD51 were considered positive if larger than 300 and 500 nm in diameter respectively. Aggregating these positive foci over both geminin+ and foci+ nuclei, and plotting their relative percentages revealed clear signal for increased RAD51 formation and thus HR competence upon ZN-c3 treatment (**Figure 4h**).

**Figure 5:**
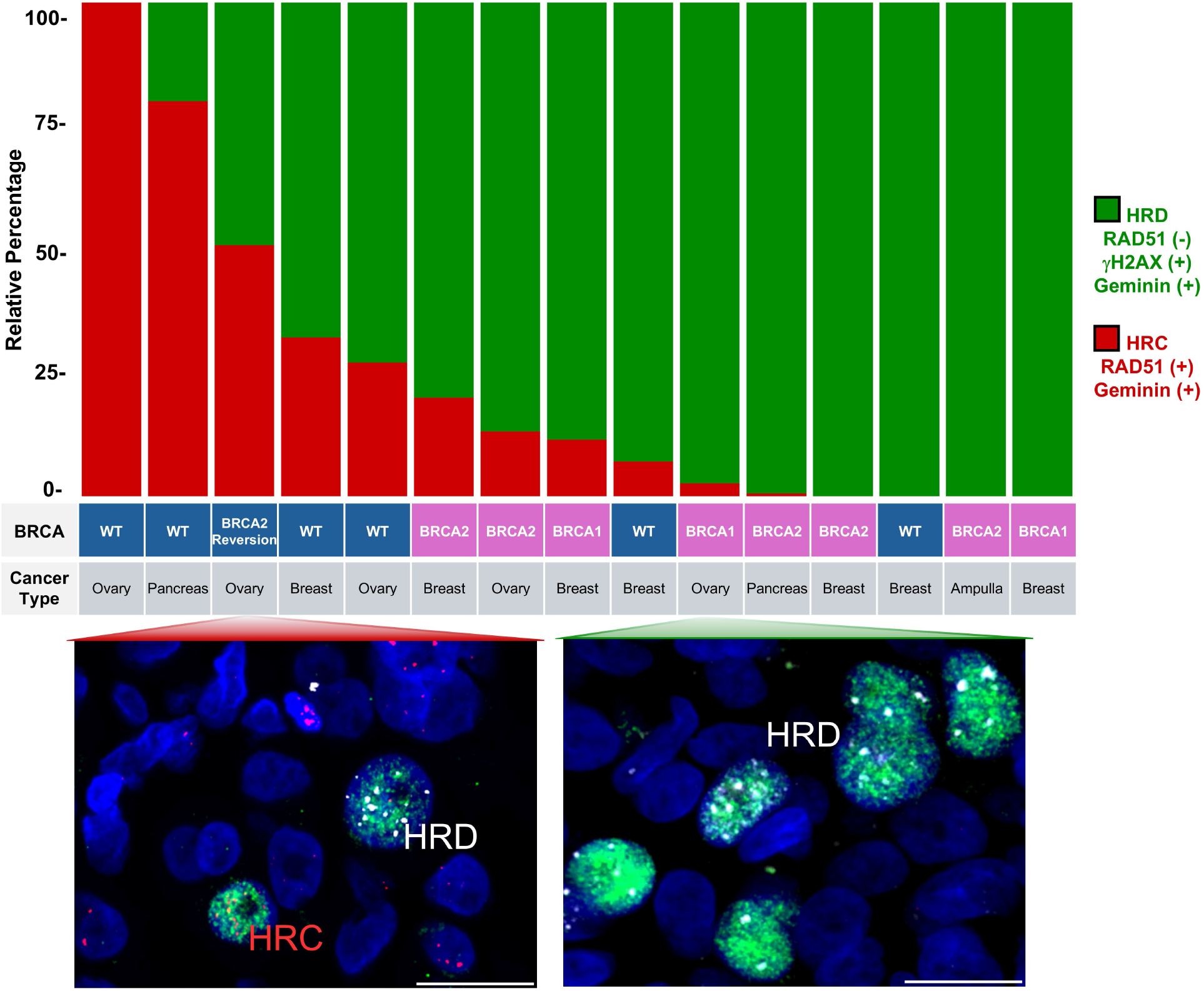
Enhancing HR Precision through Analyzing Per-Nucleus HRC/HRD Percentages. Patient stratification using HR status with varying cancer types and BRCA mutational status, patients ranked by the relative percentage of HRC/HRD nuclei after removing Uninformative nuclei. Representative images of HRC and HRD patients respectively. Scale bar shown at 10 µm.

### Functional Assay Reveals HRD Beyond BRCA Mutational Status

To demonstrate the utility of our assay in patient FFPE biopsies, we applied MAP-HR to a representative cohort of patients with diverse tumor types and varying BRCA mutational status **(Figure 5)**. Using our analysis pipeline and decision tree to categorize cell populations into HRC, HRD, and Undetermined groups, we plotted the relative percentage of HR status of each individual tumor cell in the ROI across the cohort. As the number of foci-negative G2 cells (called “Uninformative” by our decision tree) can vary substantially by patient and ROI, we removed these from visualization to better observe the relative populations of only HRC and HRD nuclei. When patients were organized by decreasing levels of HRC proportions, we found that most BRCA *wt* patients displayed high levels of HRC populations including a patient with a known BRCA2 reversion^31^ In contrast, nearly all BRCA1/2 *mt* samples displayed predominantly HRD populations. Notably, two BRCA *wt* patients showed an HRD phenotype, highlighting the importance of assessing HR deficiency with a functional assay, beyond BRCA mutational status alone to identify populations of patients likely to benefit from PARPi or platin based therapy. Our analysis of nuclei calls revealed that 5/7 BRCA *wt* patients (including the BRCA2 reversion case) exhibited an HRC phenotype, while 2/7 were HRD. Interestingly, although all 8 BRCA1/2 mutant patients had predominantly HRD populations, 3 showed a small but detectable population of HRC cells (~10-20% by relative proportion), which may contribute to the reported resistance to PARPi in some patients despite the presence of BRCA mutations^32^. Notably, in some patient slides, we observed pronounced spatial heterogeneity, with distinct clusters of HRC and HRD groups which requires further investigation to understand its implications.

## DISCUSSION

DNA-damaging therapies that induce DSBs, such as PARPi and platinum-based agents, are selectively effective in patients with HRD due to synthetic lethality^6^. Therefore, accurately identifying patients whose tumors are HRD is crucial for optimizing treatment selection and minimizing ineffectual therapies with attendant toxicity. Studies^8,11^ have shown that HRD can occur in the absence of BRCA mutations or aberrations in other members of the HR pathways, underscoring the need to detect patients with HRD profiles beyond pathway aberrations including mutation or methylation of promoters for key mediators of HR including BRCA1/2, RAD51, and PALB2^10^. Furthermore, not all patients with aberrations in the HR pathway benefit from PARPi due to healing of mutations, reversal of methylation, increased expression of hypomorphs, and inhibition of non-homologous end joining (NHEJ) through aberrations in the shieldin complex resulting in a switch to HR^33^. Thus, assessing active functional HRD through immunofluorescence-based foci assays has emerged as a valuable tool ^8,11,25^. These foci-based assays have enabled the identification of HRC and HRD status in tumors beyond those associated with BRCA1/2 mutations or other HR-related gene alterations, providing insight into the functional status of the HR pathway. However, current IF-based assays have limitations, such as the lack of both per-nucleus analysis and concurrent assessment of multi foci markers, which potentially could constrain the evaluation of HR functionality resulting in both false positive and false negative calls. Achieving precise and accurate HRD detection requires a reliable and scalable analysis platform to ensure optimal patient triage and therapeutic decision-making.

While many software solutions have successfully commercialized nuclei segmentation programs, most still lack robust intranuclear foci segmentation capabilities, highlighting the need for custom segmentation pipelines. We have thus created MAP-HR, a pipeline that obviates many of the challenges with commercial platforms. While the majority of foci-based imaging and analysis pipelines focus on 2D data, nuclei and nuclear foci are inherently 3D in nature, prompting continued research and commercial image analysis tools for their precise 3D structure^34^. 3D datasets inherently provide a better view of foci present across different focal heights, considering the typical FFPE tissue section is ~5 µm. 3D foci analysis, especially when scaled to biopsies which are cm^2^ in size, currently makes large-scale analysis on 3D foci structural details relatively infeasible. Critically, through combined orthogonal projections from 3D sections, we demonstrate how the relevant FWHM foci diameters are enhanced through axial projections, outperforming both average 2D slices and even axial foci diameters.

Importantly, these orthogonal projections found *γ*H2AX and RAD51 foci diameters to be approximately 300-500 nm respectively, which agrees with other functional studies^16,35^. Therefore, we implemented a minimum size threshold of 300 nm for *γ*H2AX and 500 nm for RAD51. While 3D datasets require slightly more input acquisition time on confocal systems, the orthogonal projection retains high-quality foci details while being much faster to analyze downstream as 2D datasets. Unlike traditional foci-detection schemes which utilize global thresholding or strictly maximal intensity positions, our pipeline extracts relevant foci segmentation based on local FWHM profiles trained via machine learning. Importantly, by extracting a wider range of foci based on shape rather than strict intensity, downstream analysis can be integrated to normalize for batch and sample specific variabilities. This alternative strategy should make a more precise and reproducible approach, especially when extending our analysis to larger clinical cohorts.

In contrast to other methods that do not assess *γ*H2AX in conjunction with RAD51 and Geminin within the same nucleus, our approach could provide a more comprehensive understanding of DNA repair mechanisms. By integrating *γ*H2AX, we mitigate the risk of inaccurately overestimating the proportion of HRD, which can occur when *γ*H2AX is not assessed in the same nucleus. Therefore, the concurrent analysis of RAD51, Geminin, and *γ*H2AX within the same nucleus on a per-nucleus basis could enhance accuracy in HR identification while also enabling the assessment of intra-tumoral heterogeneity in HR status within patient samples. Additionally, to assess HR more accurately, we removed G2 foci-negative nuclei populations (uninformative cells) from later analyses. This allows quantification which only includes cells informative for HR and performing per-nucleus analysis, our method delivers a more precise and detailed assessment of repair activity amidst DNA damage.

Importantly, through our proof-of-principle mixed BRCA *mt*/*wt* discovery cohort, we have shown the capability to apply MAP-HR pipeline to patient samples. Despite the growing use of IF-based assays in research, their clinical implementation remains severely limited. Hence, developing scalable and transparent pipelines that are accessible to the community is crucial for enhancing both accessibility and clinical adoption^36^. Considering that most groups still rely on manual counting of foci, which may introduce bias, we developed an open-source analysis pipeline that is robust and suitable for research use. Further, building on this, we plan to implement our assay in a CLIA-like environment with a focus on clinical application. MAP-HR offers a standardized and automated approach, reducing potential subjectivity and improving reproducibility in foci quantification. By extending this analysis to larger clinical cohorts, we aim to further refine the stratification of HRC/HRD levels, offering insights that correlate with treatment outcomes beyond BRCA status alone. Notably, utilizing a larger cohort from various cancer types will help determine whether a pan-cancer model or lineage-specific predictive models are more suitable for assessing HRD. Additionally, the presence of heterogeneous populations with mixed HRC and HRD clusters suggests that HR status should not be assessed in bulk. Our per-nucleus-based approach could provide valuable insights into this heterogeneity by evaluating each nucleus individually. The consequences of this heterogeneity on response to PARPi and platin will need to be assessed in future cohorts of patients with therapy outcomes. Finally, expanding the assay to include other interrelated processes, such as replication stress, could provide a more comprehensive understanding of DNA repair mechanisms and their impact on tumor biology and treatment response.

## Acknowledgements

This project was carried out with major support from the Oregon Health & Science University (OHSU) SMMART Program, Brenden Colson Center for Pancreatic Health, National Institutes of Health (NIH), National Cancer Institute (NCI), the Center for Early Detection and Advanced Research (CEDAR) and a kind gift from the Miriam and Sheldon Adelson Medical Research Fund. We would also like to thank the OHSU Knight Cancer Institute BioLibrary and Advanced Multiscale Microscopy Shared Resource for their expert technical assistance.

## Author Contributions

T.Y.O, M.J.R designed the experiments and G.B.M. supervised the project. The methodology was devised by T.Y.O. and M.J.R. who also wrote the main manuscript text. T.Y.O optimized staining and imaging protocol. T.Y.O., and G.Z. conducted the imaging experiments. M.J.R. initially coded the segmentation and feature extraction pipeline, which was troubleshooted by F.O and T.Y.O. The manuscript underwent critical review and meticulous editing by F.O and G.B.M. All authors have read and agreed to the published version of the manuscript.

---

**Supplementary Figure 1:**
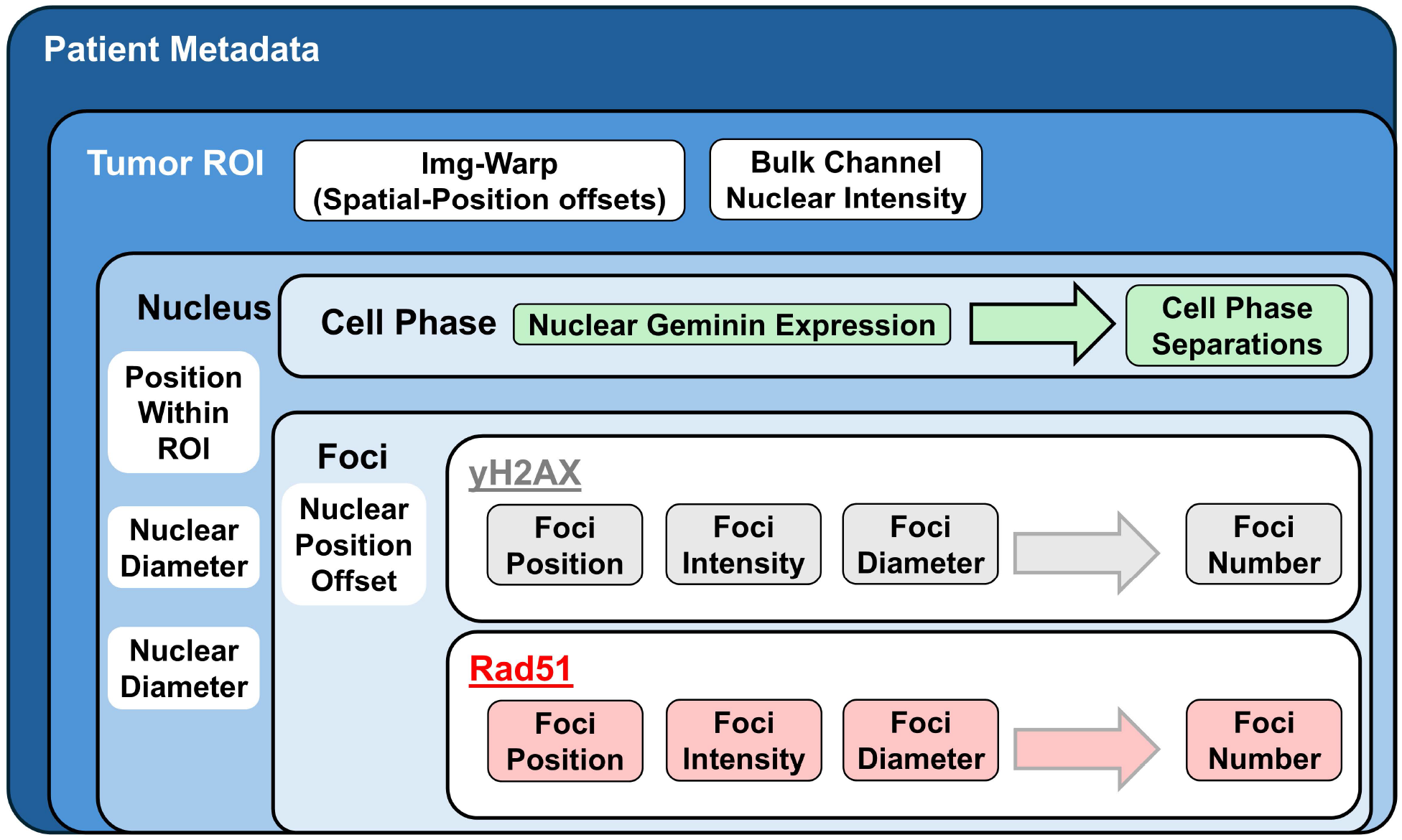
Structured Framework for Analyzing Foci-Based Nuclear Data. Idealized minimum data framework for organizing foci-based nuclear details for patient-scale analysis of HR. Data spans multiple levels from 1) Metadata, 2) Tumor-rich region annotations, 3) Nuclear features including position and cell phase via Geminin staining, 4) Nuclear positional offsets to track foci, 5) Foci-specific attributes such as position, intensity, diameter.

**Supplementary Figure 2:**
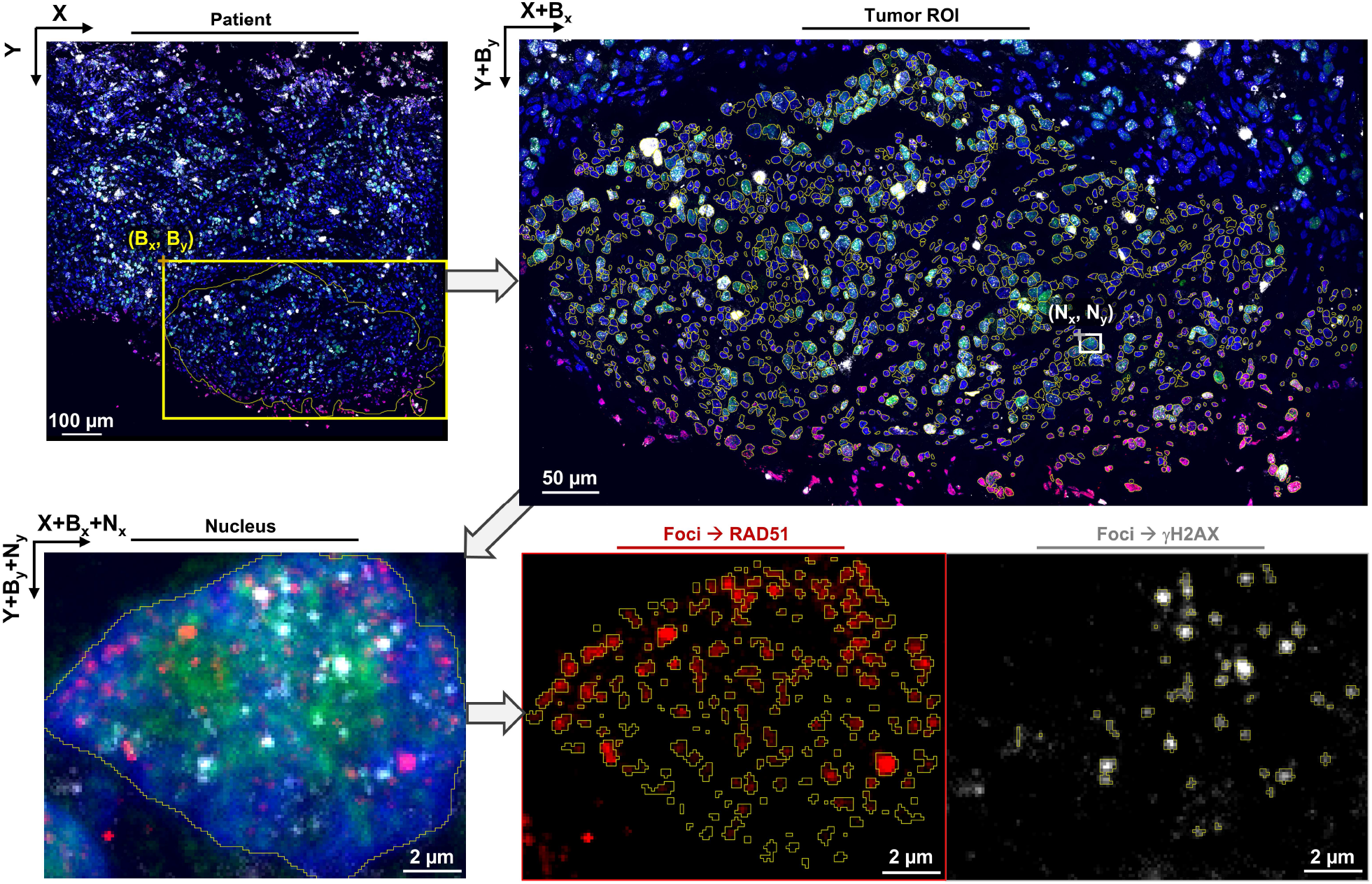
Tracking Positional Offsets through FOVs and Individual Nuclei and Matching Foci. Positional offsets needed when aggregating multi-level spatial information from FIJI. Starting from original image with X,Y global coordinates, tumor-rich regions of interest (ROI) are selected for nuclear segmentation. These result in new bounding boxes (B_x_, B_y_) specific to each ROI, which has its own inherently global offset needed to correct for the new 0,0 starting position for each ROI on the upper left. This can be denoted as needing to re-align the ROI-specific coordinates back onto the original full patient image using X_global_=X+B_x_ and Y_global_=Y+B_y_. This corrects nuclear coordinates back to the original image. Similarly, at the foci-level, the nuclear bounding box (N_x_, N_y_) specific to each nucleus ROI. Finally, foci coordinates in the original patient image can be corrected as: using X_global_=X+B_x_+N_x_ and Y_global_=Y+B_y_+N_y_.

**Supplementary Figure 3:**
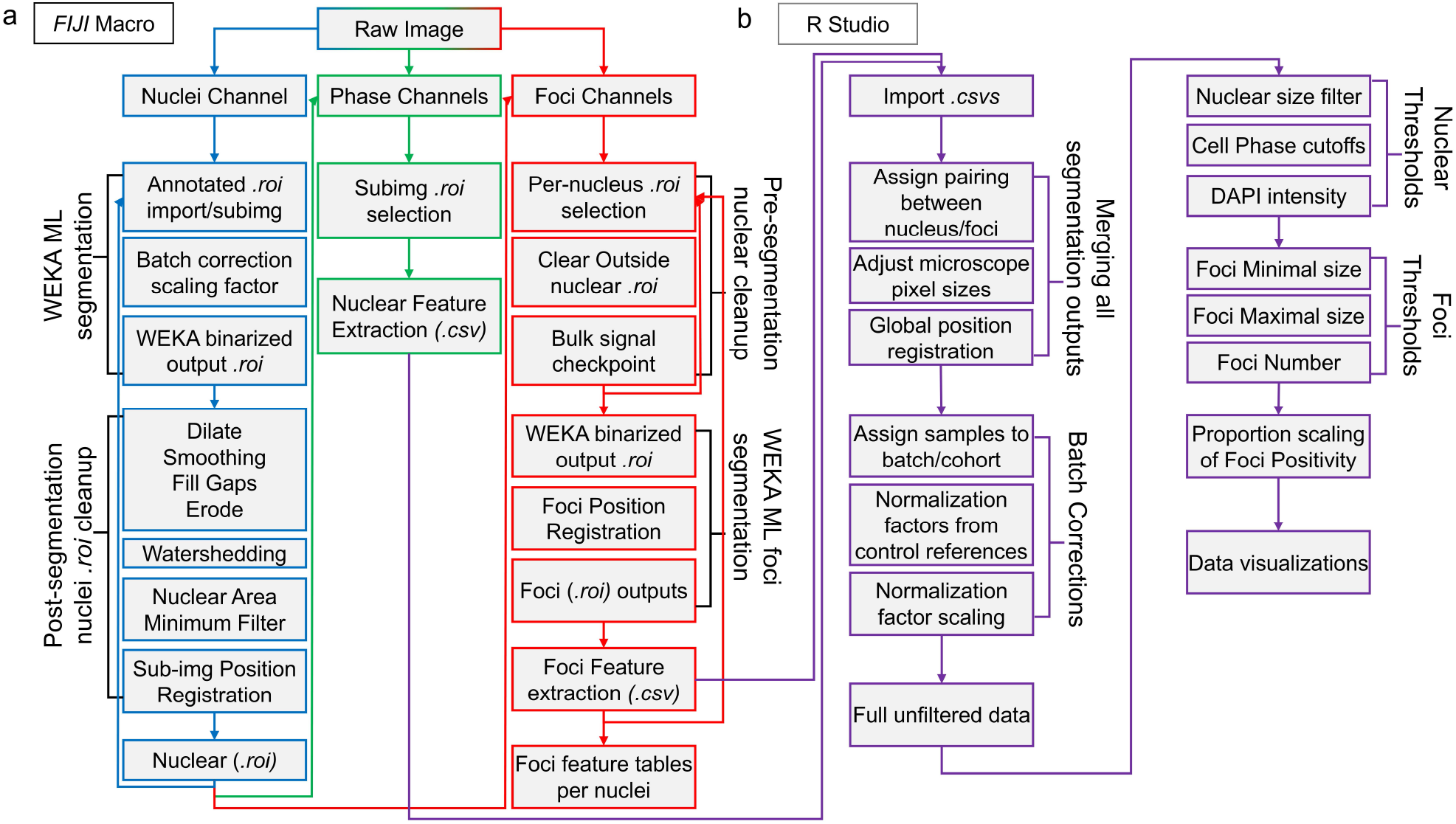
MAP-HR Overall Feature Extraction Workflow and Downstream Analysis in R. MAP-HR overall workflow comprised of both: *a. FIJI* Macro workflow for segmentation and feature extraction for cell phase and foci level details. b. Downstream R processing for data organization, positional alignment, normalization, thresholding, and visualizations.

**Supplementary Figure 4:**
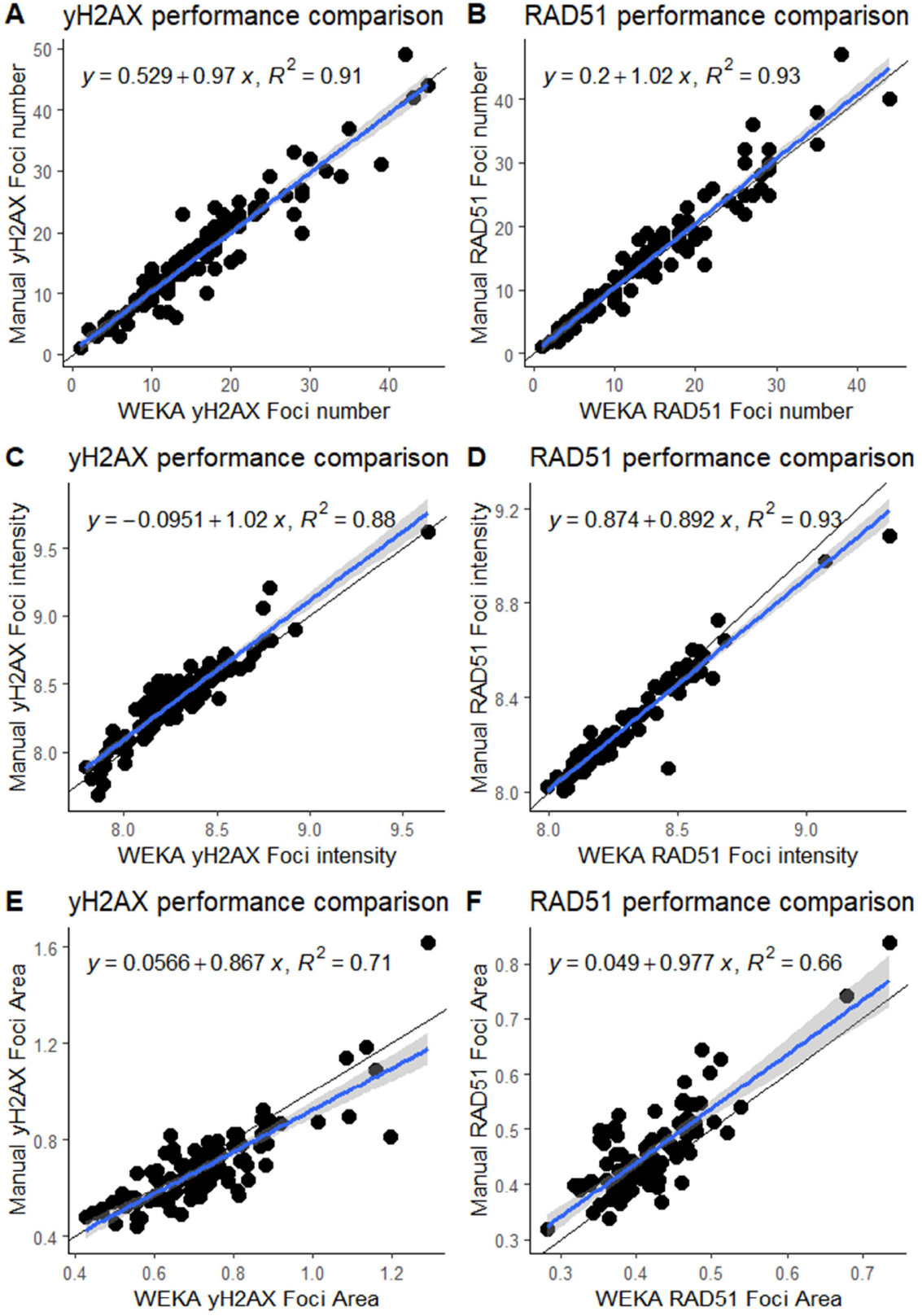
Evaluating Performance of Foci Segmentation Features Comparing Manual vs WEKA. Foci segmentation performance correlations from 100 representative nuclei comparing Manual vs WEKA outputs as measured by: a) yH2AX Foci number per nucleus b) RAD51 Foci number per nucleus c) Average yH2AX Pixel intensity per nucleus d) Average RAD51 Pixel intensity per nucleus e) Average yH2AX Foci Area per nucleus f) Average RAD51 Foci Area per nucleus

**Supplementary Figure 5:**
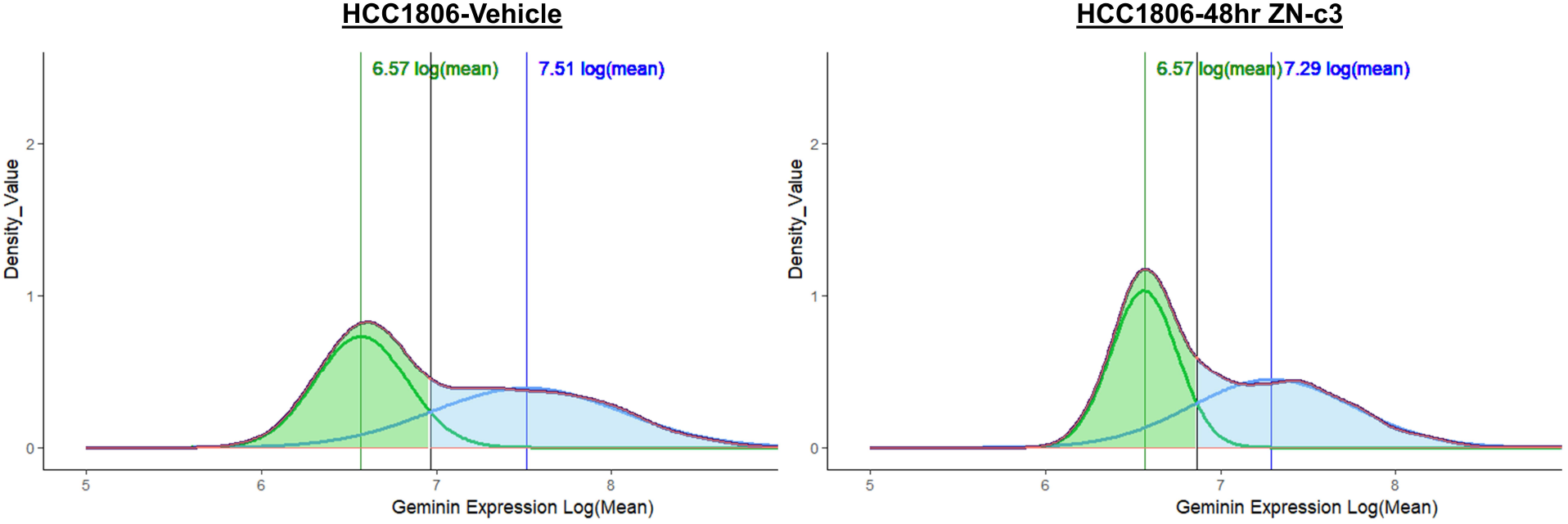
Gaussian Mixture Modelling for G2 Phase Determination. Gaussian mixture models for geminin G2 cutoffs as shown for HCC1806-Vehicle control vs ZN-c3 treated. Intercept between both G1 and G2 population peaks were used to define geminin positivity. Each fit gaussian shown with colored vertical line indicating its center intensity value on Log scaling. Analyses were performed in *R* and plotted using *ggplot2* package using *geom_density*.

