## Supplemental Figures 1-5 for "Automated Machine Learning Profiling with MAP-HR for Quantifying Homologous Recombination Foci in Patient Samples"

**This PDF file includes:**

Supplementary Figures 1 to 5

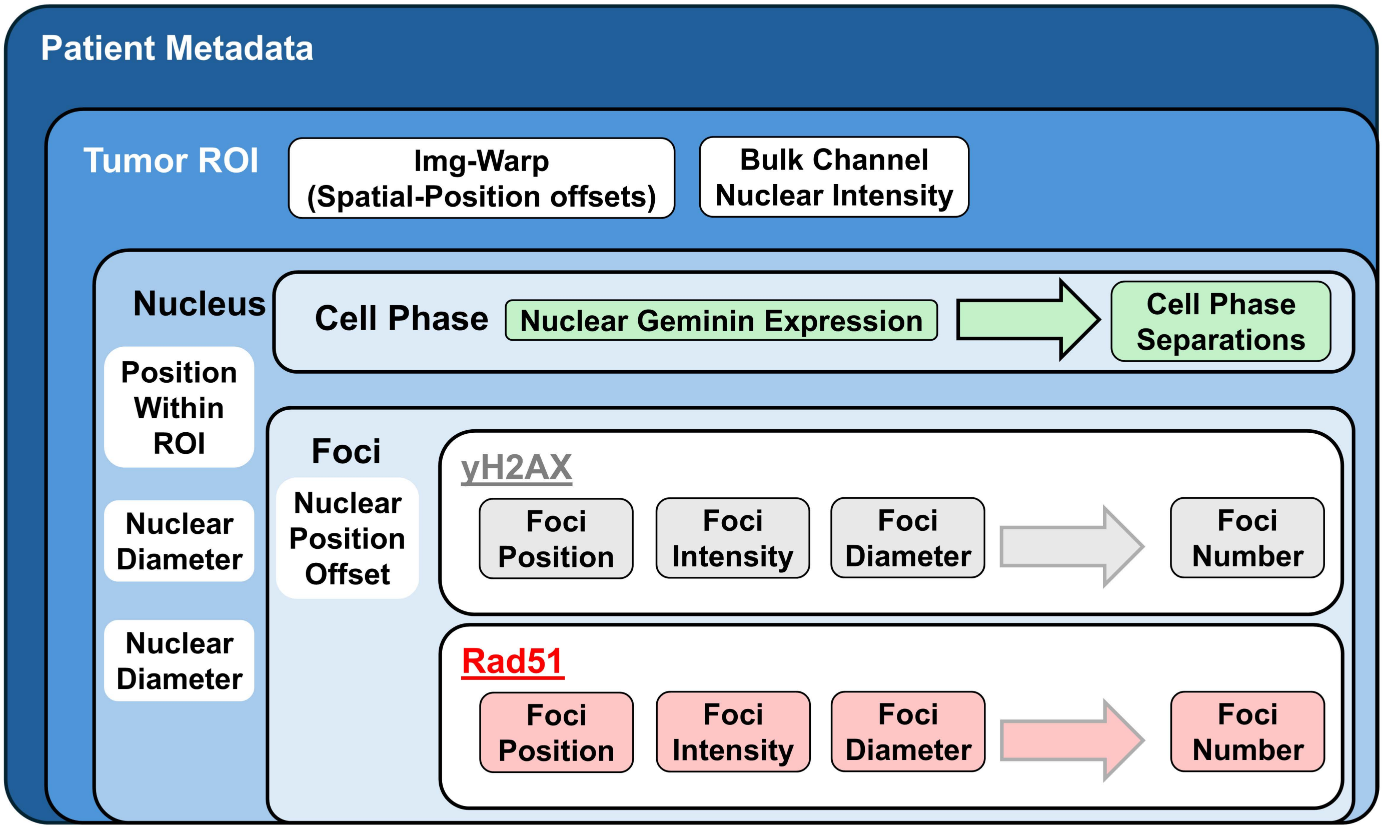


**Supplemental Figure 1: Structured Framework for Analyzing Foci-Based Nuclear Data**

Idealized minimum data framework for organizing foci-based nuclear details for patient-scale analysis of HR. Data spans multiple levels from 1) Metadata, 2) Tumor-rich region annotations, 3) Nuclear features including position and cell phase via Geminin staining, 4) Nuclear positional offsets to track foci, 5) Foci-specific attributes such as position, intensity, diameter.

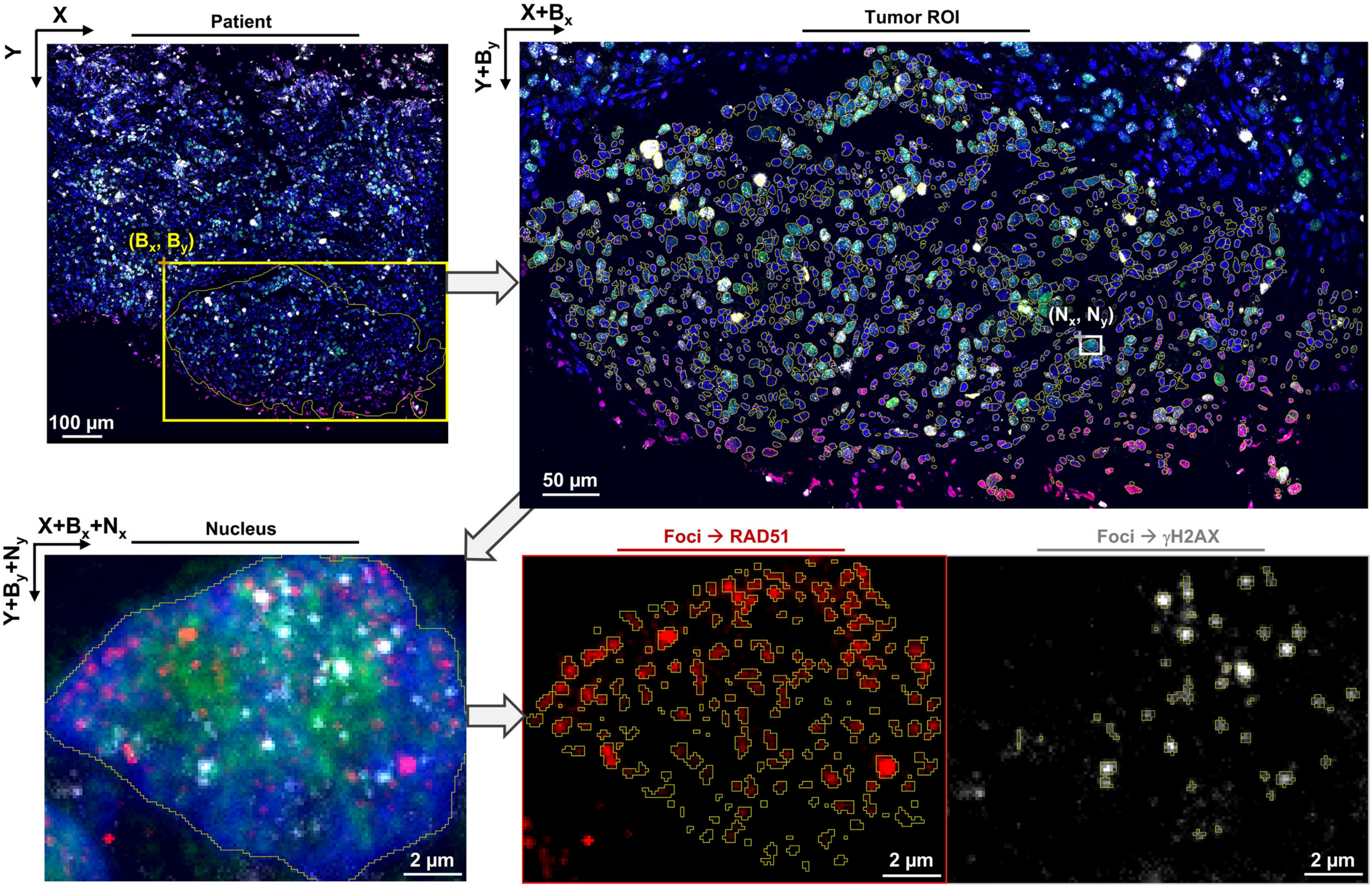


**Supplemental Figure 2: Tracking Positional Offsets through FOVs and Individual Nuclei and Matching Foci**

Positional offsets needed when aggregating multi-level spatial information from FIJI. Starting from original image with X,Y global coordinates, tumor-rich regions of interest (ROI) are selected for nuclear segmentation. These result in new bounding boxes ($B_x$, $B_y$) specific to each ROI, which has its own inherently global offset needed to correct for the new 0,0 starting position for each ROI on the upper left. This can be denoted as needing to re-align the ROI-specific coordinates back onto the original full patient image using $X_{global}=X+B_x$ and $Y_{global}=Y+B_y$. This corrects nuclear coordinates back to the original image. Similarly, at the foci-level, the nuclear bounding box ($N_x$, $N_y$) specific to each nucleus ROI. Finally, foci coordinates in the original patient image can be corrected as: using $X_{global}=X+B_x+N_x$ and $Y_{global}=Y+B_y+N_y$.

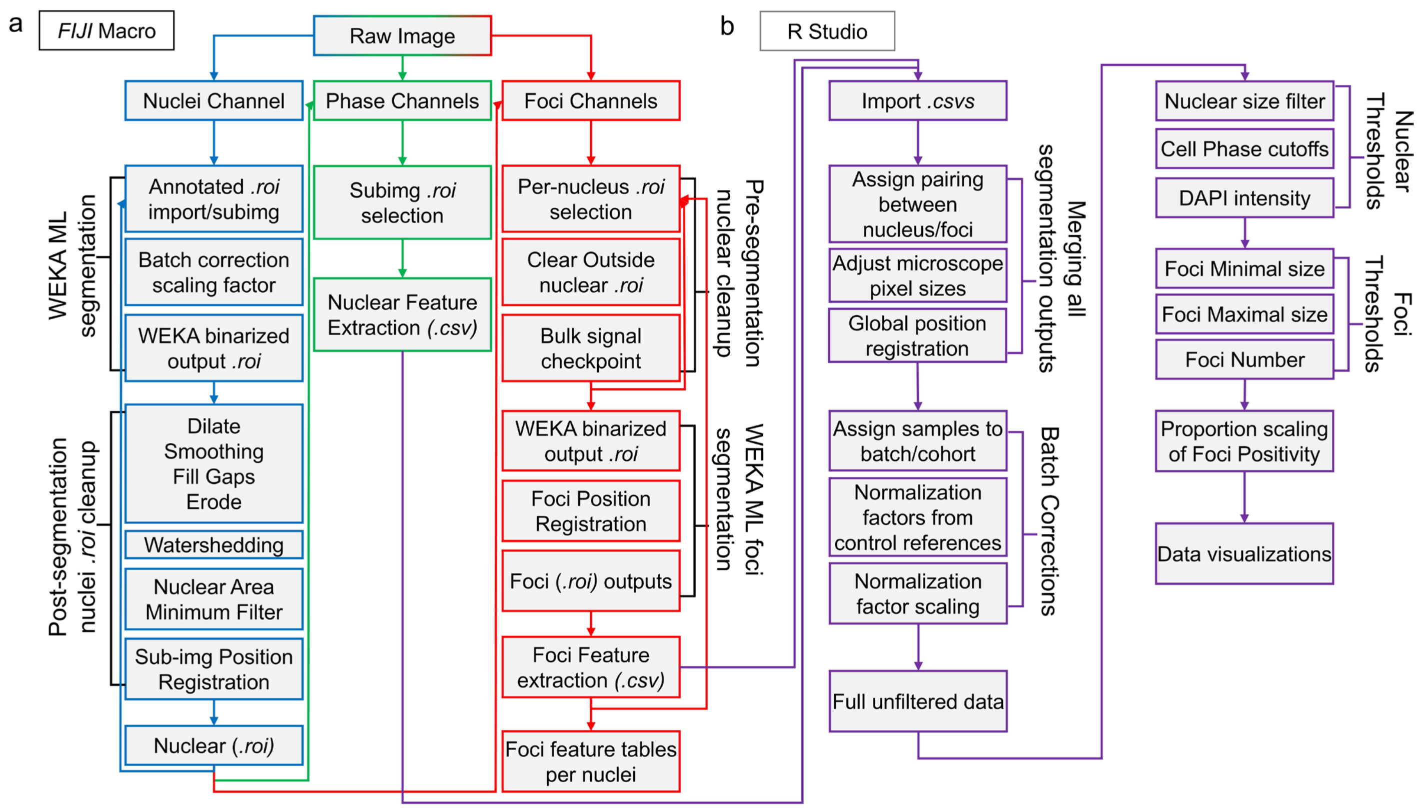


**Supplemental Figure 3: MAP-HR Overall Feature Extraction Workflow and Downstream Analysis in R**

MAP-HR overall workflow comprised of both:

a. *FIJI* Macro workflow for segmentation and feature extraction for cell phase and foci level details.

b. Downstream R processing for data organization, positional alignment, normalization, thresholding, and visualizations.

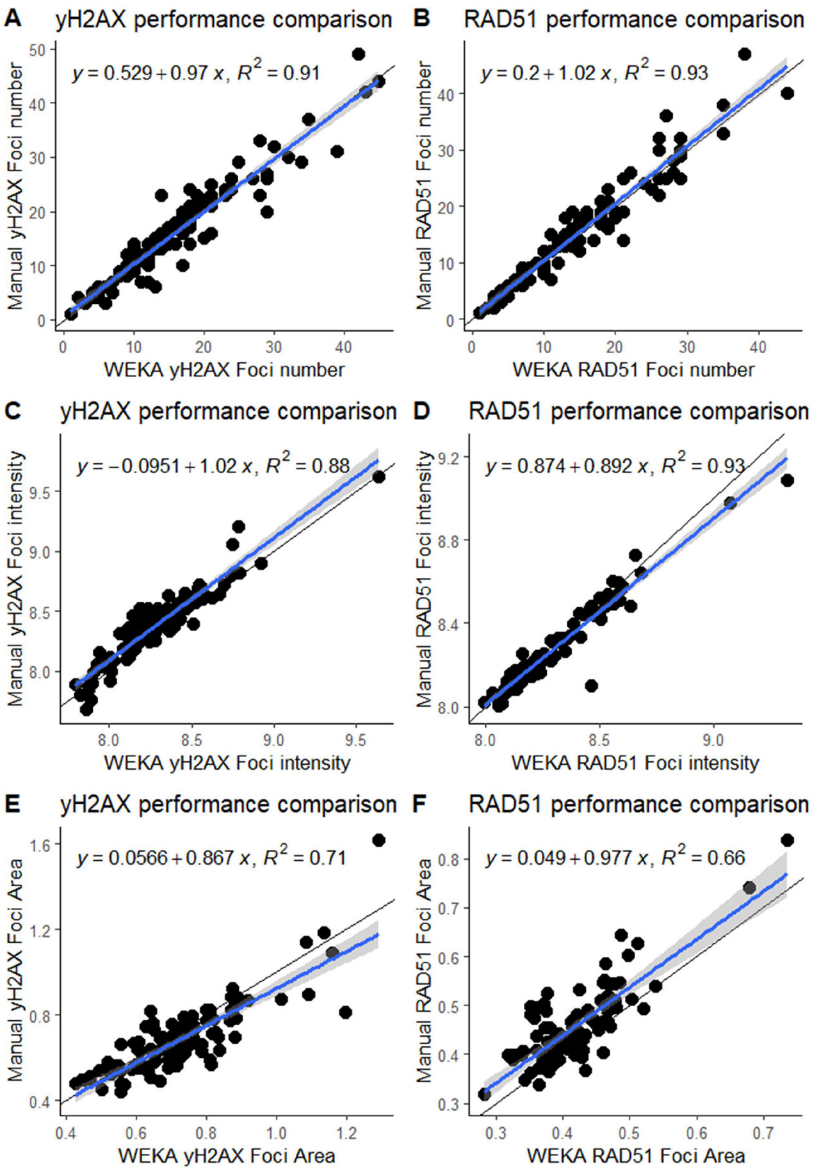


**Supplemental Figure 4: Evaluating Performance of Foci Segmentation Features Comparing Manual vs WEKA**

Foci segmentation performance correlations from 100 representative nuclei comparing Manual vs WEKA outputs as measured by:

a) yH2AX Foci number per nucleus
b) RAD51 Foci number per nucleus
c) Average yH2AX Pixel intensity per nucleus
d) Average RAD51 Pixel intensity per nucleus
e) Average yH2AX Foci Area per nucleus
f) Average RAD51 Foci Area per nucleus

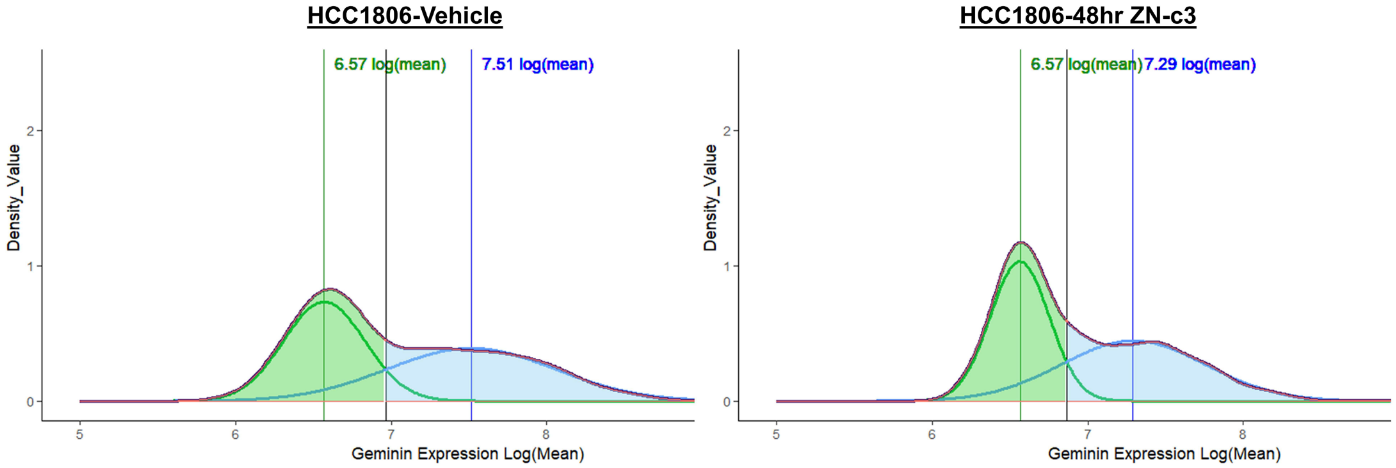


**Supplemental Figure 5: Gaussian Mixture Modelling for G2 Phase Determination**

Gaussian mixture models for geminin G2 cutoffs as shown for HCC1806-Vehicle control vs ZN-c3 treated. Intercept between both G1 and G2 population peaks were used to define geminin positivity. Each fit gaussian shown with colored vertical line indicating its center intensity value on Log scaling. Analyses were performed in *R* and plotted using *ggplot2* package using *geom_density*.
